# Multi-Species Phenotypic Screening across Disease Models of Mucolipidosis Type IV

**DOI:** 10.1101/2021.03.05.434120

**Authors:** Andrea Hadjikyriacou, Sangeetha Iyer, Joshua D. Mast, Nina DiPrimio, John Concannon, Joshua Ketterman, Frederic Sigoillot, Tamy P. Rodriguez, Feba S. Sam, Hillary Tsang, Madeleine Prangley, Julide Bilen, Kausalya Murthy, Tom A. Hartl, Christophe Antczak, Jeremy L Jenkins, Nathan T. Ross, Beat Nyfeler, Rishi K. Jain, John A. Tallarico, Ethan O. Perlstein, Stephen M. Canham

## Abstract

Invertebrate model organisms (the nematode *Caenorhabditis elegans* and the fruit fly *Drosophila melanogaster*) are valuable tools to bridge the gap between traditional in vitro discovery and preclinical animal models. Invertebrate model organisms are poised to serve as better disease models than 2D cellular monocultures for drug discovery, as well as easier and more cost-effective to scale up than 3D organoids/assembloids or co-cultures. A strength of model organisms is the opportunity to probe conserved biology such as lysosomal function and autophagy in a physiological setting. However, invertebrate models are not without pharmacokinetic and pharmacodynamic challenges, such as poor tissue penetration and confidence in a compound’s mechanism of action. To confront those challenges, we took advantage of the Novartis mechanism-of-action box (MoA Box), a compound library of well-annotated and drug-like chemical probes. Curious as to how the MoA Box, comprised of chemical probes optimized for mammalian targets, would fare in an invertebrate setting we screened the MoA Box across three different models of the lysosomal storage disease mucolipidosis Type IV (MLIV). MLIV is caused by mutations in the lysosomal transient receptor potential ion channel mucolipin-1 (TRPML1) resulting in hyper-acidic lysosomes and dysregulated autophagy. The overlap of screening hits from worm, fly, and patient fibroblast screens identified cyclin-dependent kinase (CDK) inhibition as an evolutionarily conserved disease modifier and potential drug repurposing strategy.

**Summary statement:** A trio of phenotypic screens across *Drosophila*, *C. elegans,* and *H. sapiens* models of mucolipidosis IV was performed and identified overlapping hits including cyclin-dependent kinase inhibitors.

## Introduction

Animal models are an integral part of the drug discovery process for the development of new medicines. For decades, the pharmaceutical industry has used rodents, dogs, and non-human primates to test the effects of drug compounds before advancing to clinical trials in humans. In contrast, the application of *Drosophila melanogaster* (fly), *Caenorhabditis elegans* (worm), and *Danio rerio* (zebrafish) within pharmaceutical research has often been overlooked despite full sequencing of their genomes in the 1990s, tools to enable genetic manipulation, and the substantial body of basic research to understand underlying pathways of biology and diseases. The short history of *Drosophila*, *C. elegans*, and *Danio rerio* at pharmaceutical companies such as Exelixis (Artavanis-Tsakonas, 2004), Bristol Meyer Squibb (Carroll et al, 2003), Novartis (Fishman, 2012; Ready, 2002), and others have largely focused on functional genomics applications that include: 1) the use of invertebrate organisms to model disease phenotypes and better understand the fundamental underpinnings of biology (Chang et al, 2008; Singh et al, 2019; Tang et al, 2020); 2) genetic screens for the potential identification of new therapeutic targets (Chen et al, 2008; Mahoney et al, 2006); and 3) chemical genetic approaches towards identifying the target or mechanism of action of small molecule drug candidates (FitzGerald et al, 2006; Lackner et al, 2005).

Despite the greater translatability and higher yields of unbiased phenotypic screening, often resulting in first-in-class drugs, pharmaceutical companies have hesitated to invest in high-throughput screening with invertebrate animal models despite their low cost and demonstrated success in academia (Mofatt, et al., 2017; Zheng, et al., 2013; Haasen, et al., 2017; Swinney & Anthony, 2011; Eder, et al., 2014; Segalat, 2007; O’Reilly, et al., 2014; Peterson & Fishman, 2011) (Segalat, 2007). A significant concern is the predictive value of invertebrate model organisms in drug discovery, already a known liability of the ubiquitously used mouse model system, which has 95-97% genetic similarity to humans (Pound & Ritskes-Hoitinga, 2018; Lander, et al., 2001). Animal-human species differences are an appreciable concern. However, the *Drosophila* genome is 60% homologous to that of humans, less redundant, and approximately 75% of the genes responsible for human diseases have homologs in flies. (Ugur, et al., 2016). The *C. elegan*s genome has over 50% genetic homology to humans and a much smaller degree of splicing than what occurs in humans, resulting in a simplified proteome (Kaletta & Hengartner, 2006; Kim, et al., 2007). These genetic similarities and simplifications have the potential to enable discoveries of fundamental disease modifier pathway biology. Early studies at Novartis using 27 small molecules targeting seven signaling pathways demonstrated a high degree of conserved compound activity between *Drosophila* and vertebrates (Bangi, et al., 2011). That early observation inspired us to expand the multi-species approach (Strynatka, et al., 2018) and perform phenotypic screens across two invertebrate model organisms and a patient-derived fibroblast cell line. We decided to use the Novartis Mechanism-of-Action Box (MoA Box), which in total is a collection of 4,185 well-defined chemical probes for >2,100 mammalian targets (Canham, et al., 2020). We hoped that the MoA Box would be well-suited to overcome known *Drosophila* and *C. elegans* xenobiotic defenses that ordinarily render many pharmacological tools ineffective. The compounds within the MoA Box have high cellular permeability, potency, and selectivity with drug-like properties. The collective MoA Box was analyzed by an *in silico* model of bioaccumulation in *C. elegans* where ~25% of the MoA box compounds score >1 and thus likely to accumulate in worms. (Burns, et al., 2010) (**Supplemental Table 1A**).

In our phenotypic screens, we chose to probe lysosomal function as it is well conserved across species and offered an attractive approach to exploring autophagy in a whole-animal setting (Hansen, et al., 2018). Additionally, there is a plethora of genetically engineered invertebrate model organisms for lysosomal storage disorders validated and reported in the literature (de Voer, et al., 2008; Levine & Klionsky, 2004; Zhang & Peterson, 2020). The endolysosomal ion channel, transient receptor potential channel mucolipin-1 (TRPML1), has garnered recent interest in drug discovery as its role as a key regulator of lysosomal function has evolved (Chen, et al., 2014; Shen, et al., 2012). Mucolipin-1 functions to maintain calcium, zinc, and iron homeostasis; regulates lysosomal acidification and autophagy; and helps drive lysosomal maturation and biogenesis (Venkatachalam, et al., 2015; Lloyd-Evans & Platt, 2011; Cheng, et al., 2010; Zeevi, et al., 2007).

Mucolipidosis type IV (MLIV) is a lysosomal storage disease caused by mutations in the *MCOLN1* gene that encodes TRPML1. MLIV disease results in developmental delay with observed motor, cognitive, and visual impairments. Despite a 19-year effort to study the natural history of MLIV disease (NCT00015782, NCT01067742), the limited number of MLIV patients has hindered understanding of MLIV disease, and to date there are no effective treatments (Wakabayashi, et al., 2011). We therefore hoped that our multi-species approach leveraging *Drosophila*, *elegans,* and patient fibroblasts might yield novel drug targets or druggable pathways for MLIV disease.

## Results

### Characterization and establishment of phenotype in invertebrate and fibroblast models of MLIV disease

We took advantage of the previously published invertebrate MLIV models created in *Drosophila* and *C. elegans*. The *Drosophila* genome encodes one homolog, aptly named *trpml*, with ~40% amino acid similarity to human TRPML1-3. Similar to the human disease MLIV, deficiency of *trpml* in flies cause neurodegeneration, motor impairments, and pupal lethality. (Venkatachalam, et al., 2008) *Trpml*^*−/−*^ homozygote animals can be rescued by consuming a high protein diet during the larval stage, enabling pupae to fully eclose and hatch into adult animals (Wong, et al., 2012). Akin to MLIV patients, *trpml-*deficient flies exhibit defects in autophagy and motility, demonstrated by the fly’s inability to climb. We generated *trpml*^*−/−*^ homozygotes through a 2-kilobase deletion by CRISPR/Cas9 editing, resulting in a *trpml* null animal that displayed pupal lethality as demonstrated by the failure of pupae to eclose (**SI Figure 1.1**). Both the *trpml*^*+/−*^ heterozygote flies and *trpml*^*−/−*^ flies reproduced phenotypes previously reported. Therefore, we devised a phenotypic screen to rescue the *trpml*^*−/−*^ homozygotes to adulthood, which we would measure by video capture of movement of eclosed adult flies in 96-well plates (**Figure 1A**, **SI Figure 1.2**, **SI Video 1.3**, & **SI Figure 1.4**).

**Figure 1:**
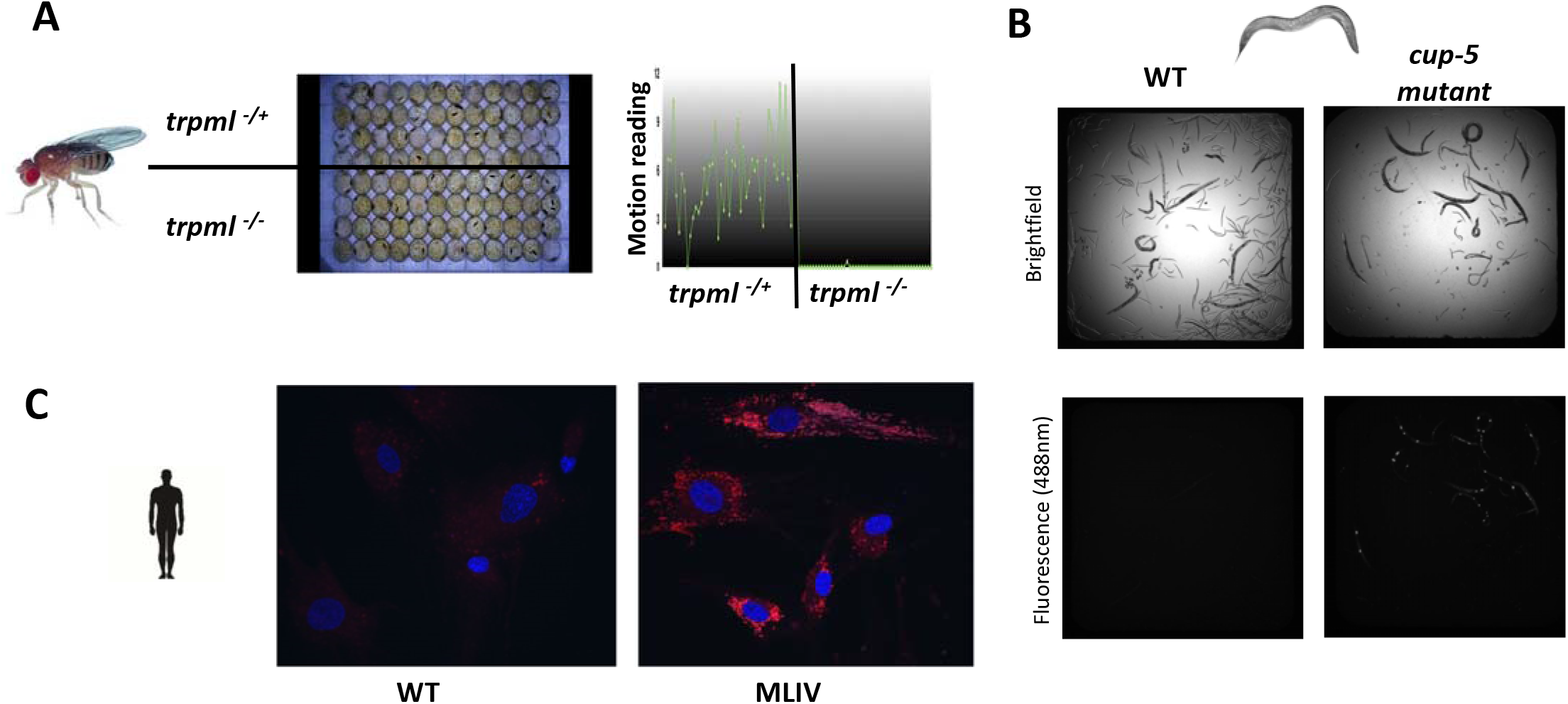
A) *Trpml* fly development and locomotion assay. Normal development to pupal stage was observed on both *trpml*^*−/+*^ and *trpml*^*−/−*^ animals. However only *trpml*^*−/+*^ animals develop into adults where their activity is measured by video recordings of movements at day 13, 14, and 15 after plating the 1^st^ instar larvae. B) Phenotypic comparison of wild-type (WT) and *cup-5* mutant worms. Five days after plating L1 larvae, the *cup-5* mutant worms have smaller brood size and under fluorescent imaging can observe green fluorescent protein (GFP) accumulation in coelomocytes. C) A comparison of wild-type (WT) and MLIV patient fibroblasts displays marked accumulation of LysoTracker (red) in acidic vesicles of the MLIV fibroblast cells; DAPI (blue).

The *C. elegans* genome encodes a homolog of TRPML1 called *cup-5* (coelomocyte uptake defect-5), in which autophagy is defective. A previously published *cup-5* mutant worm displays embryonic lethality and autophagy defects manifested by the accumulation of green fluorescent protein (GFP) in specialized phagocytic cells called coelomocytes (Fares & Greenwald, 2001; Schaheen, et al., 2006; Hersh, et al., 2002). An attractive feature to the *cup-5* mutant model is that treatment with the small molecule methyl pyruvate demonstrated partial rescue of the embryonic lethal phenotype (Schaheen, et al., 2006). From the Fares Lab we acquired the *C. elegans cup-5* hypomorphic strain *ar465[Pmyo-3::ssGFP],* caused by a G401E missense mutation which exhibits characteristic GFP accumulation in coelomocytes, along with wild-type *arIs37[Pmyo-3::ssGFP]* expressing wild-type CUP-5. We were able to replicate the reported phenotype (**Figure 1B** & **SI Figure 1.5**) (Fares & Greenwald, 2001).

To enable comparison of animal-human species differences in our chemical genetic screen we needed to develop a human cell-based assay. In MLIV patient fibroblast cells, lysosomes are enlarged, hyper-acidified, and exhibit accumulation of phospholipids and mucopolysaccharides when compared to non-disease cells. We first obtained four MLIV patient fibroblast cell lines from the Coriell Institute cell repository. Upon initial characterization, we found that the GM02527 line, containing a homozygous IVS3-2A>G insertion, grows quickly and is well suited for a high-throughput, image-based phenotypic screen. The IVS3-2A>G insertion causes a loss of reading frame, skipping exon 4 and is found in 72% of the MLIV patient population (Sun, et al., 2000). To establish a cellular phenotype, we evaluated an unbiased approach through a screen of the Fluopack collection of fluorescently labeled metabolites and analytes (Kang, et al., 2020). Additionally, we evaluated a targeted approach examining fluorescent and immunofluorescent markers such as LysoTracker (lysosomal accumulation and pH), FluoZin (zinc accumulation), LC3 (autophagy), p62 (autophagy), filipin (cholesterol accumulation) and BODIPY LacCer (sphingolipid accumulation) (Vergarajauregui, et al., 2008). While LysoTracker, LC3 and p62 displayed differential accumulation when comparing wild-type vs MLIV patient fibroblasts (**SI Figure 1.6**), only the LysoTracker cellular phenotype could be rescued upon treatment with the TRPML1 channel activator, MK6-83, and did not require starvation (**SI Figure 1.7**) (Chen, et al., 2014). We therefore proceeded to use the LysoTracker readout as our primary assay (**Figure 1C**) as it enabled the inclusion of positive control compounds and we would use p62 and LC3 as secondary assays.

### Phenotypic screening across models of MLIV disease

To conduct our *trpml Drosophila* chemical genetic screen, MoA box compounds were plated at a concentration of 20μM with the food source into 96-well plates. Homozygote and heterozygote fly larvae were then dispensed and incubated for 13-14 days. The plates were processed using a custom-built fly imager and the motion from flies in the wells was quantified. Plates were also analyzed manually and scored for partial and full emergence of adults at day 15 (**SI Figure 2.1**). Comparison of positive control *trpml*^*−/+*^ heterozygous flies across plates from the negative control DMSO-treated *trpml*^*−/−*^ homozygous flies showed strong separation of the data in both the automated activity assay as well as through the manual scoring method (**SI Figure 2.2**). The MoA box collection was screened in triplicate with the *trpml*^*−/−*^ flies and upon characterization with the automated activity assay there were few hits that replicated across experiments, with only three hits replicating in two experiments and 46 hits occurring from any replicate. Luckily the manual scoring format provide a z′-score of >1.5 with 41 hits that replicated in more than two experiments (**SI Figure 2.3**). The low hit discovery rate in our fly screen may not be surprising as drug exposure related artifacts such as reduced feeding relating to the preference of flies for each compound, uncertainty about the amount of food consumed, compound stability in fly food, stochasticity due to small numbers of animals per well, and tissue concentration achieved by each compound are known (Qi, et al., 2015; Ja, et al., 2007; Marx, 2015; Iyer, et al., 2019). To enable our ability to compare the fly hits with the orthogonal screening efforts in worms and fibroblasts, we chose to be more inclusive of our hits for analysis. We combined the automated activity scored hits along with the manually scored hits and to bring the total number to 97 fly hits (**Figure 2A**, **SI Table 1B**).

**Figure 2:**
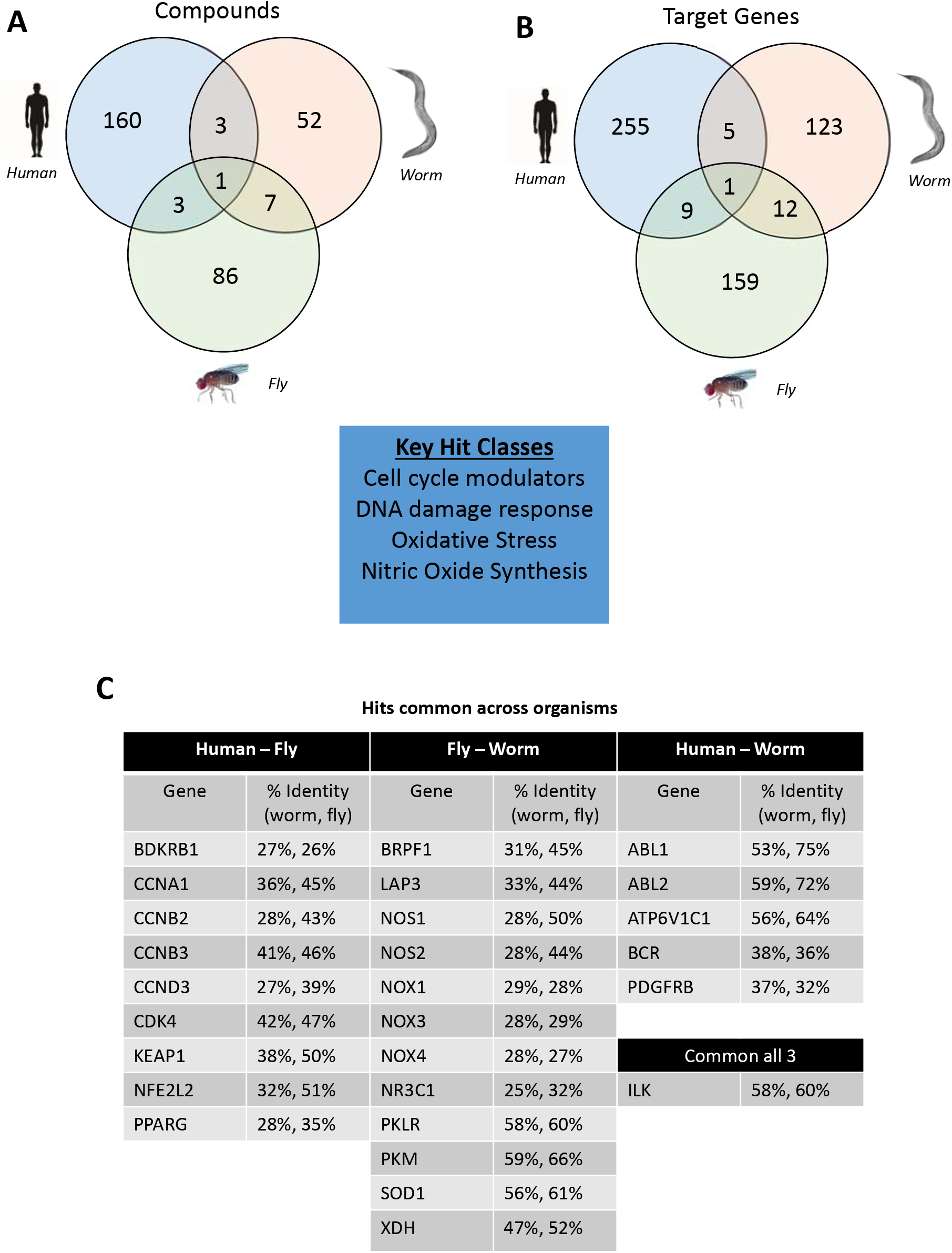
A) Venn diagram displaying overlapping number of compound hits across human (blue), worm (orange), and fly (green) screens using the MoA Box library. B) Venn diagram showing the corresponding target genes from the compound hits that overlap across the human, worm, and fly MoA Box screens. Key hit classes/groupings of the drug targets (blue box), and include cell cycle modulators, DNA damage response, oxidative stress, and nitric oxide synthesis pathways. C) Table of drug targets that were common in the MoA Box screens across Human– Fly screens, Fly–Worm screens, Human–Worm screens, and those that were common in all three screens. Table shows Gene as the human gene code found in UniProt and the percent similar identity of the human gene for the closest BLAST hit in worms and flies.

In parallel, the MoA box was screened with the *C. elegans cup-5* mutant. To perform this image-based screen, synchronized L1 larvae were dispensed in 96-well plates along with negative (DMSO-treated) and positive (CUP-5 wild-type *C. elegans*) controls. The *cup-5* screen was performed with compounds at a final concentration of 25μM dispensed in liquid broth in quadruplicate to ensure reproducibility. After incubation at 20°C for five days, the worms were chemically immobilized to facilitate imaging (**SI Figure 2.4**). Image analysis calculated the area per well occupied by worms as well as the GFP signal per well to establish a GFP signal/worm area ratio for each individual well. Positive and negative controls in the screen displayed strong separation across all plates (**SI Figure 2.5**). The primary assay readout we chose was a ratio of GFP to area occupied by worms, but several additional sub-categories of signals could also be identified: a) diffuse GFP fluorescence that was not concentrated in punctae b) complete clearance of punctate GFP fluorescence c) GFP fluorescent punctae that are smaller in size and intensity than their respective controls (**SI Figure 2.6**). Overall, the *cup-5* worm screen yielded 64 hits that reproduced in over three replicates, with an overall hit rate of just over 2% (**Figure 2A**, **SI Table 1C** & **SI Table 2**). Surprisingly, this hit rate was similarly low when compared with other phenotypic *C. elegans* screens despite the use of the MoA Box (Burns, et al., 2006; Kwok, et al., 2006; Iyer, et al., 2019). The higher *cup-5* worm hit rate than the *trpml* fly screen (1.45%) might be explained by the worms being continuously soaked in the compound laden media to foster compound uptake and bioaccumulation. Interestingly, the hits identified from the *C. elegans* screen had reasonable correlation with their *in silico* predicted scores for compound bioaccumulation in *C. elegans* when compared to the entire content of the MoA Box (vida supra). In total, 28 or 44% of the hits scored >1 and thus over 50% likely to bioaccumulate in the *C. elegans.* (**SI Table 1C**) (Burns, et al., 2010).

To evaluate the translatability of the invertebrate model screens, we ran the MoA Box screen in MLIV patient fibroblasts using LysoTracker as the imaging readout. Fibroblast cells were seeded into 1536-well plates and were incubated with compounds plated in 8-point dose response, in duplicate. After 48 hours, the plates were imaged by live cell imaging and fluorescence signal quantified (**SI Figure 2.7**). As a positive control, we switched from our initial control of MK6-83 to bafilomycin A1. Bafilomycin A1, which causes lysosomal de-acidification by inhibiting the vacuolar H^+^-ATPase and prevents accumulation of LysoTracker into acidic vesicles, had greater reproducibility than MK6-83 (Yamamoto, et al., 1998). Overall, the fibroblast screen showed good separation of positive and negative controls across all plates (**SI Figure 2.8**). Hits were identified as greater than two standard deviations from the negative control and an activity (AC_50_), determined by Cell Profiler signal metric Granularity2, of less than 10μM (Stoter, et al., 2019). We identified MoA box hits annotated for ATP6V1C1, the same target as the positive control bafilomycin A1, giving further confidence to the screen design. After removing cytotoxic compounds determined as having a low cell count from the DAPI nuclear staining, we identified 167 hits from the LysoTracker screen. (**Figure 2A** & **SI Table 1D**). It is hard to determine, but we speculate that the higher observed hit rate of 6% in the fibroblast screen may be attributed to the nature of the indirect and simplified LysoTracker assay readout rather than other potential factors in contrast to the counterpart screens in model organisms.

### Comparison of screening hits across species and hit validation

Plotting the screening hits across the disease models of mucolipidosis type IV demonstrates positive overlap across the three species-specific assays (**Figure 2A**). In total, there are 14 compounds that scored as a hit in at least two assays, and only one compound that was active in all three assays. As compounds may have more than one annotated target, i.e., polypharmacology, the compounds were converted to their corresponding annotated gene targets to arrive at overlapping gene targets of the three systems (**Figure 2B**). Altogether, 27 genes were correlated with compounds active in more than two assays. A BLAST gene sequence comparison of the mammalian gene target with the *Drosophila* or *C. elegans* homologs did not identify a higher degree of similarity, as what might be presumed. To expedite validation of the screens, we focused on the human MLIV fibroblasts. Confirmed gene hits could be rapidly validated in the fibroblast assay because it is more reproducible, more amenable to testing in dose-response, and tractable for rapid turn-around genetic manipulation. The 27 gene targets annotated for our overlapping hits were expanded to an additional compound set of ~100 compounds similarly annotated for the 27 overlapping gene targets, and then profiled in the LysoTracker fibroblast assay. The expansion set provided additional confidence in the positive gene target and to increase confidence in gene targets initially only identified as overlapping in the fly and worm screens. Our expansion effort narrowed the focus to approximately 26 genes that included cyclin-dependent kinases (CDKs), integrin-linked protein kinase (ILK), and c-Abl gene targets.

### Inhibition of CDKs rescues disease phenotype of MLIV and enhance autophagy

The intersection of our multi-species phenotypic screens enabled us to focus on a few drug target gene hypotheses. In order to prioritize hits, we ran our top compounds in the LysoTracker fibroblast assay. After conducting live cell imaging with LysoTracker, the cells were fixed and immunostained for p62 in order to demonstrate that the lack of LysoTracker accumulation was not an artifact of inhibiting autophagy or the lysosomotropic properties of the test compounds (Ashoor, et al., 2013; Lu, et al., 2017). Lead molecules consisting of CDK inhibitors palbociclib, ribociclib, a non-lysosomotropic ribociclib analog named Cpd-2 (Fassl, et al., 2020), BIM-1 (Davis, et al., 1992b; Davis, et al., 1992a), and CHEMBL3685712 (Zhang, et al., 2004), exhibited suppression of LysoTracker accumulation without the accumulation of p62 (**Figure 3A** & **Figure 3B**). To evaluate if autophagy was enhanced, LC3B immunoblotting was conducted to evaluate autophagic flux in MLIV fibroblasts under starvation conditions. Fibroblasts were treated with compounds and starved for 24 and 48 hours; lysates were then collected for immunoblot analysis for LC3B protein processing. Ribociclib and BIM-1 induced increased LC3-II processing, as shown by immunoblotting and the quantification ratios of LC3-I and LC3-II, indicating increased autophagosome formation (**Figure 4A** & **Figure 4B**). Since LC3-I and LC3-II levels on immunoblots can sometimes differ due to antibody reactivity (Mizushima & Yoshimori, 2007), for a better indicator of autophagosome formation we calculated the ratio of LC3-II/(LC3-I + LC3-II) using densitometry analysis of the blots in ImageJ, and compared it to the fold increase over DMSO-treated starvation conditions. In 24 hour starvation conditions, ribociclib and BIM-1 induced the highest fold change over the control (approximately 4-fold, **Figure 4B**, top panel); CHEMBL3685712 induced a two-fold increase; Cpd-2 showed no change. After 48 hours of treatment and starvation, we observe a higher fold increase following treatment with ribociclib and BIM-1 (approximately 15-fold increase, and can be seen with disappearing LC3-I band in the immunoblot) (**Figure 4B**, bottom panel). From those results, we can interpret that CDK inhibition enhances autophagosome accumulation; however, it does not imply enhanced autophagic degradation, or flux. After validating ILK and CDKs as gene targets from our screen with mechanistically targeted chemical probes, we next sought to obtain orthogonal genetic validation.

**Figure 3:**
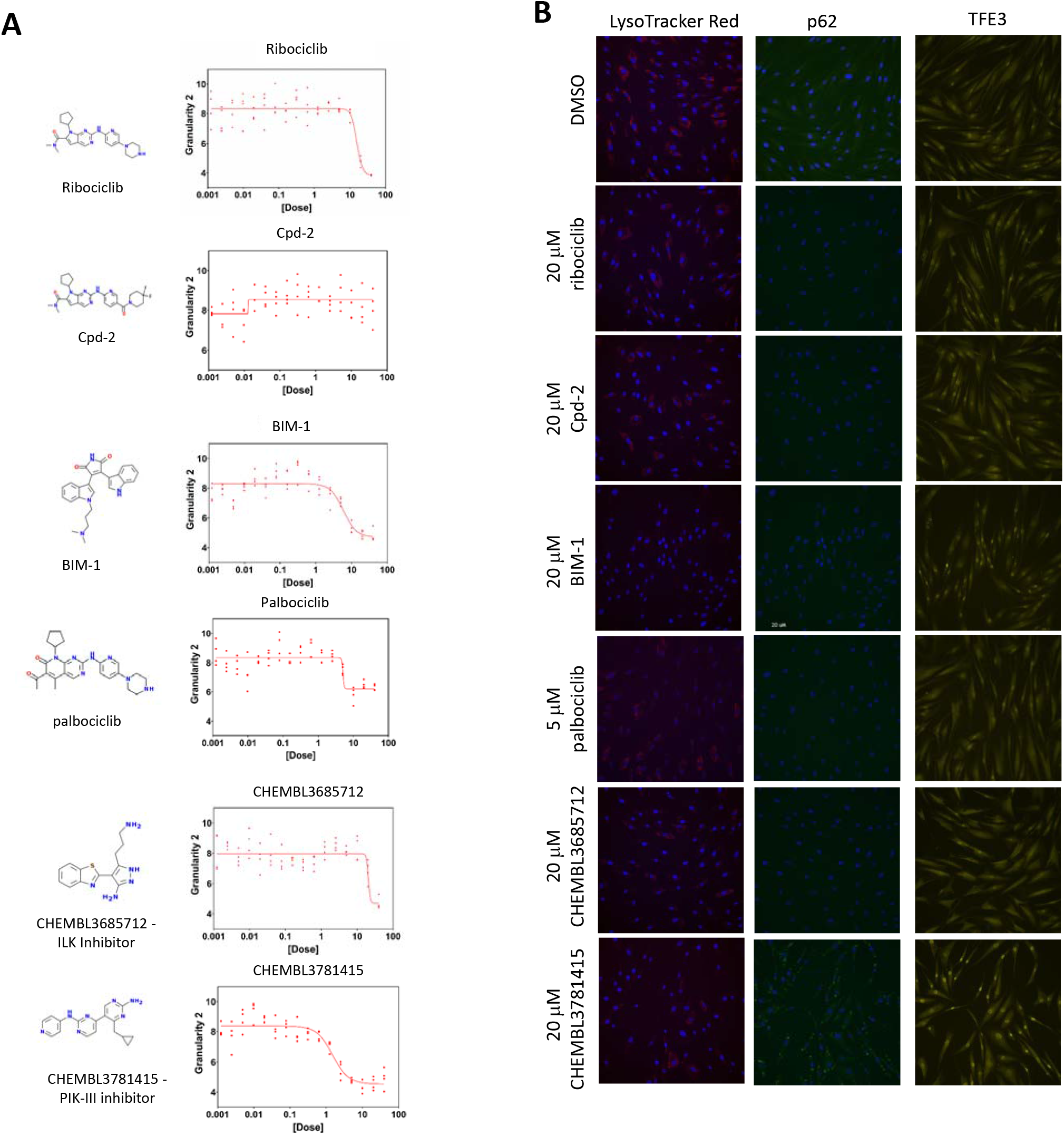
A) Compound structures and dose response curves for best five validated compounds (ribociclib, Cpd-2, BIM-1, palbociclib, and CHEMBL3685712) and one control (CHEMBL3781415 – PIK-III inhibitor). Dose response curves depict quantified LysoTracker signal metric “Granularity 2” (y-axis) as a function of dose concentration (μM; x-axis). Each dose shows four replicate points of quantified LysoTracker signal acquired from high content images that were analyzed using Cell Profiler for analysis. B) Representative images for LysoTracker (Red, left panel), p62 (green, middle panel), and TFE3 (yellow, right panel) staining after treatment with the indicated pharmacological inhibitors (20μM), except palbociclib (5μM).

**Figure 4:**
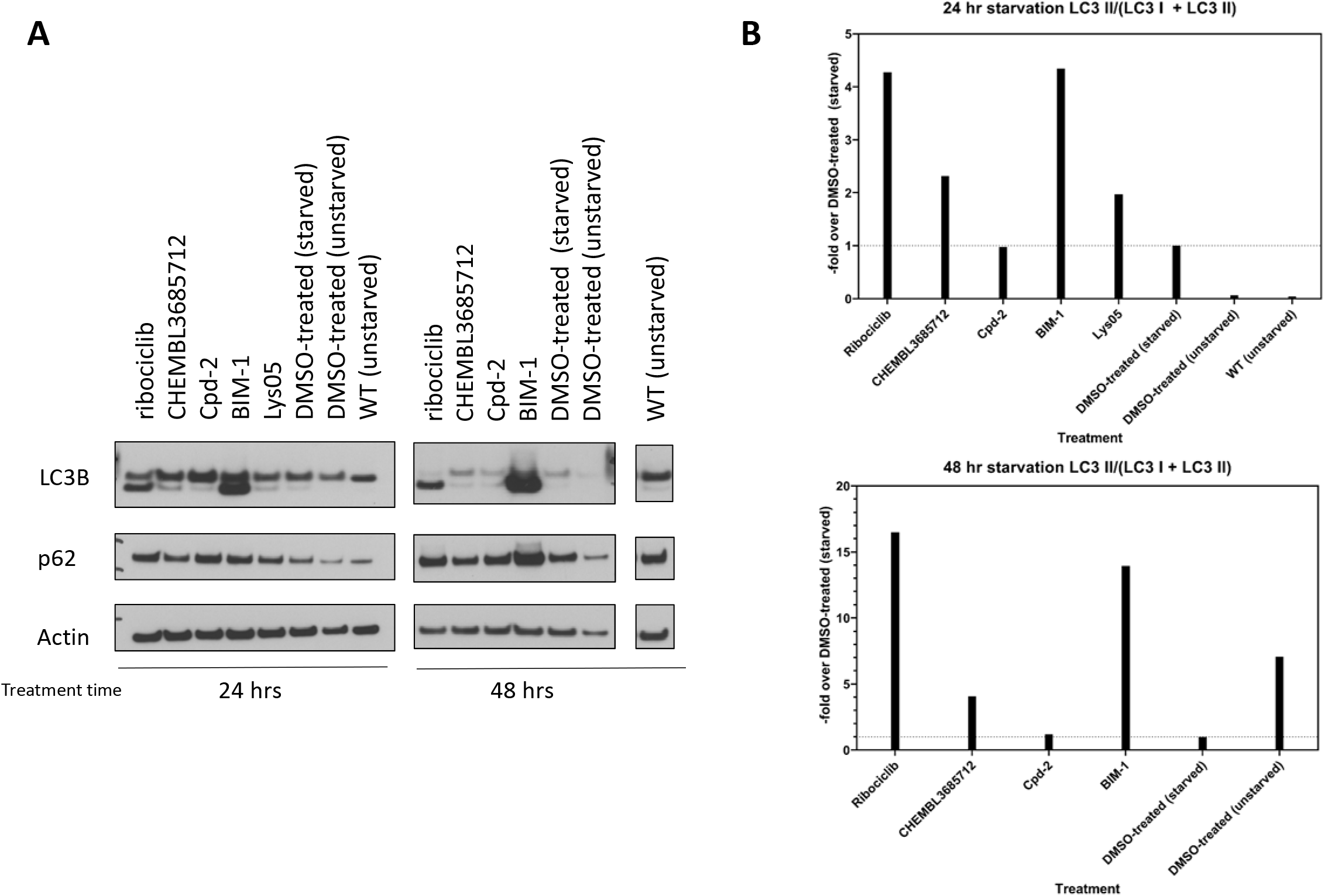
A) Immunoblots of 0.1% serum starved MLIV fibroblast cells treated with the indicated compounds at 20μM (Lys05 was treated at 1μM) for 24 or 48 hours as indicated. Cell lysates were prepared and 5 μg were loaded on SDS-PAGE gel for protein separation and then blotted for LC3B, p62, and actin. B) Immunoblot densitometry analysis of LC3-I (top) and LC3-II (lower) bands conducted with ImageJ to determine the total area under the curve and reported as a ratio of LC3-II / (LC3-I + LC3-II) and normalized to DMSO-treated cells under starved conditions.

Because kinase selectivity with small molecule inhibitors can be challenging, we decided to validate the top gene target candidates at the mRNA level by siRNA screening of the top candidates in the LysoTracker fibroblast assay as well as by LC3B immunoblotting. MLIV fibroblasts were nucleofected with siRNA ON-TARGETplus SMARTpool for the various CDKs (CDK2, CDK4, CDK5, CDK6, CDK9), ILK, GBA, and non-targeting siRNA as a control, and grown for either 72 hours in 384-well plates for imaging with LysoTracker or for five days in 6-well dishes for immunoblotting. Separating the lysates on SDS-PAGE and running an immunoblot for LC3B, we observed low induction of LC3-II (**Figure 5A**), with only CDK6 showing an approximately two - fold increase in LC3-II/(LC3-I + LC3-II) ratio (**Figure 5B**). TFE3 immunostaining showed slight increase in nuclear localization in CDK4, CDK5 and CDK6 siRNA-treated conditions compared to a non-targeting control (**Figure 5C**). LysoTracker imaging was inconclusive, perhaps because electroporation induces cells stress and may induce lysosome exocytosis (Napotnik, et al., 2016; Thompson, et al., 2014).

**Figure 5:**
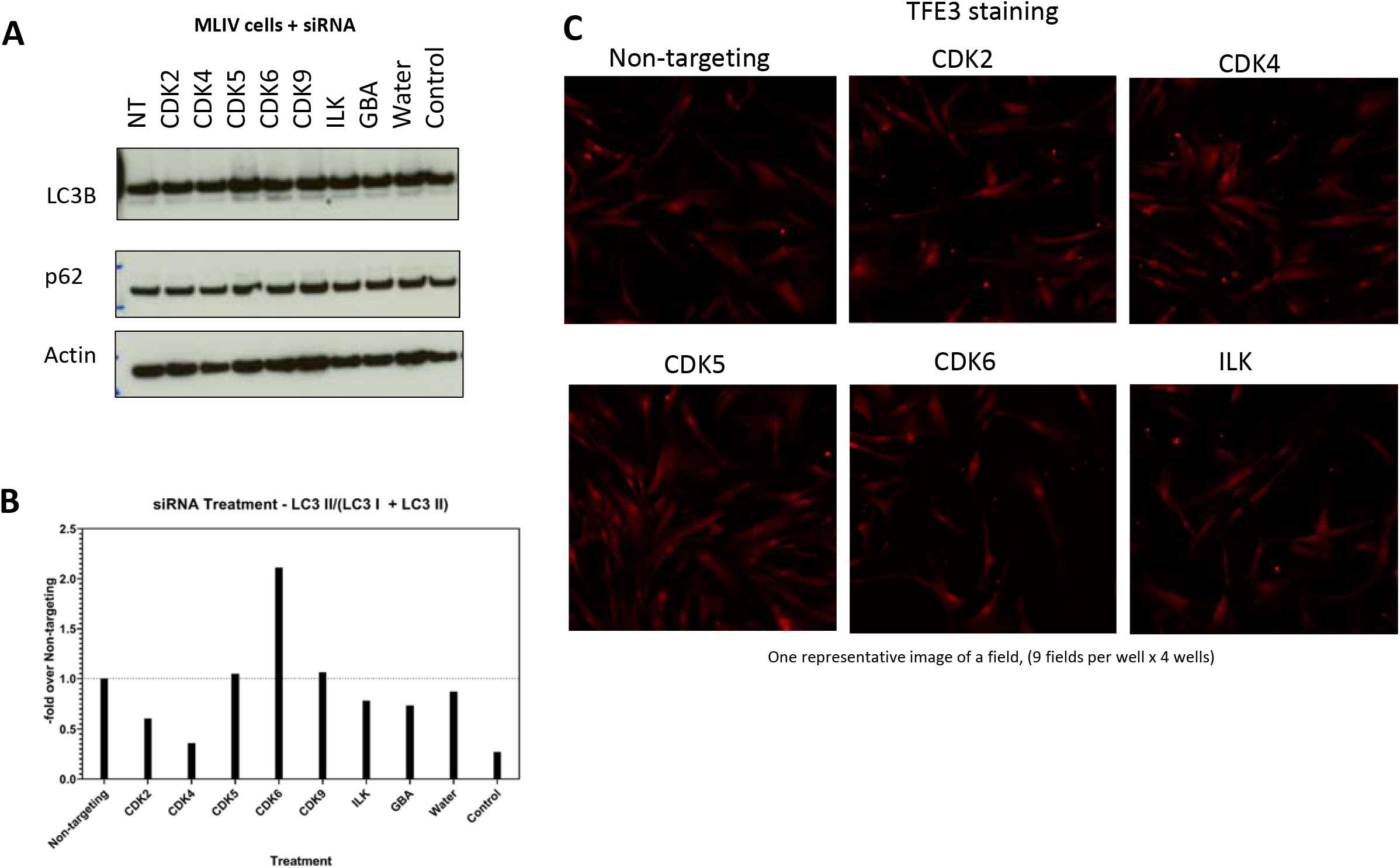
A) Immunoblots of MLIV fibroblast cells treated with siRNA pools for various gene targets (CDK2, CDK4, CDK5, CDK6, CDK9, ILK, GBA, non-targeting (NT)) for five days. Cell lysates were prepared and 5 μg were loaded on SDS-PAGE gel for protein separation and then blotted for LC3B, p62, and actin. Water denotes MLIV fibroblast cells that were only subjected to electroporation conditions and Control denotes MLIV fibroblast cells that were not subjected to treatment with siRNA or electroporation conditions. B) Immunoblot densitometry of LC3-I (top) and LC3-II (lower) bands analysis from immunoblot was performed with ImageJ software. Analysis was conducted to determine the total area under the curve and obtain a ratio of LC3-II / (LC3-I + LC3-II), normalized to DMSO-treated cells under starved conditions. Water denotes MLIV fibroblast cells that were only subjected to electroporation conditions and Control denotes MLIV fibroblast cells that were not subjected to treatment with siRNA or electroporation conditions. C) Representative images of TFE3 staining of representative siRNA-treated MLIV fibroblast cells (non-targeting, CDK2, CDK4, CDK5, CDK6 and ILK).

We have demonstrated that CDK4/6 inhibition could improve lysosomal biogenesis and autophagic flux. However, we felt it important to demonstrate that CDK4/6 inhibition restores the balance of lysosomal calcium levels, signifying a correction or bypass of the faulty TRPML1 Ca^2+^ channel. To probe lysosomal calcium levels in MLIV fibroblast cells, we used CalipHlour_*Ly*_, a pH-correctable, DNA-based fluorescent reporter (Narayanaswamy, et al., 2019). We treated MLIV fibroblasts with five compounds (ribociclib, Cmpd-2, BIM-1, CHEMBL3685712, Lys05), incubated them for 48 hours, and then added CalipHlour_*Ly*_ nine hours prior to live cell imaging (McAfee, et al., 2012). Images were processed in order to determine the pH in individual lysosomes, and to apply correction factors to compute the value of lysosomal calcium ions with single lysosome resolution, using images generated in different channels – Green, Orange and Red. When comparing wild-type (WT) and DMSO-treated MLIV cells we observed a difference in their pH and [Ca^2+^] maps due to dysregulation of lysosomal pH and calcium levels in MLIV cells (**Figure 6**, top panels). Unfortunately, cells treated with ribociclib and BIM-1 were not analyzable because of probe mislocalization. This observation may result from the intrinsic lysosomotropic property of ribociclib and BIM-1, which disrupts the pH and localization of the CalipHlour_*Ly*_ probe. Alternatively, these compounds could be enhancing autophagic flux (**Figure 4B**), and inducing rapid degradation of the probe. Upon treatment with Cpd-2, a non-lysosomotropic CDK4/6 inhibitor, we observed a return to WT levels, which was not observed with the other test compounds, CHEMBL3685712 and control compound Lys05 (**Figure 6**, bottom panels) (McAfee, et al., 2012).

**Figure 6:**
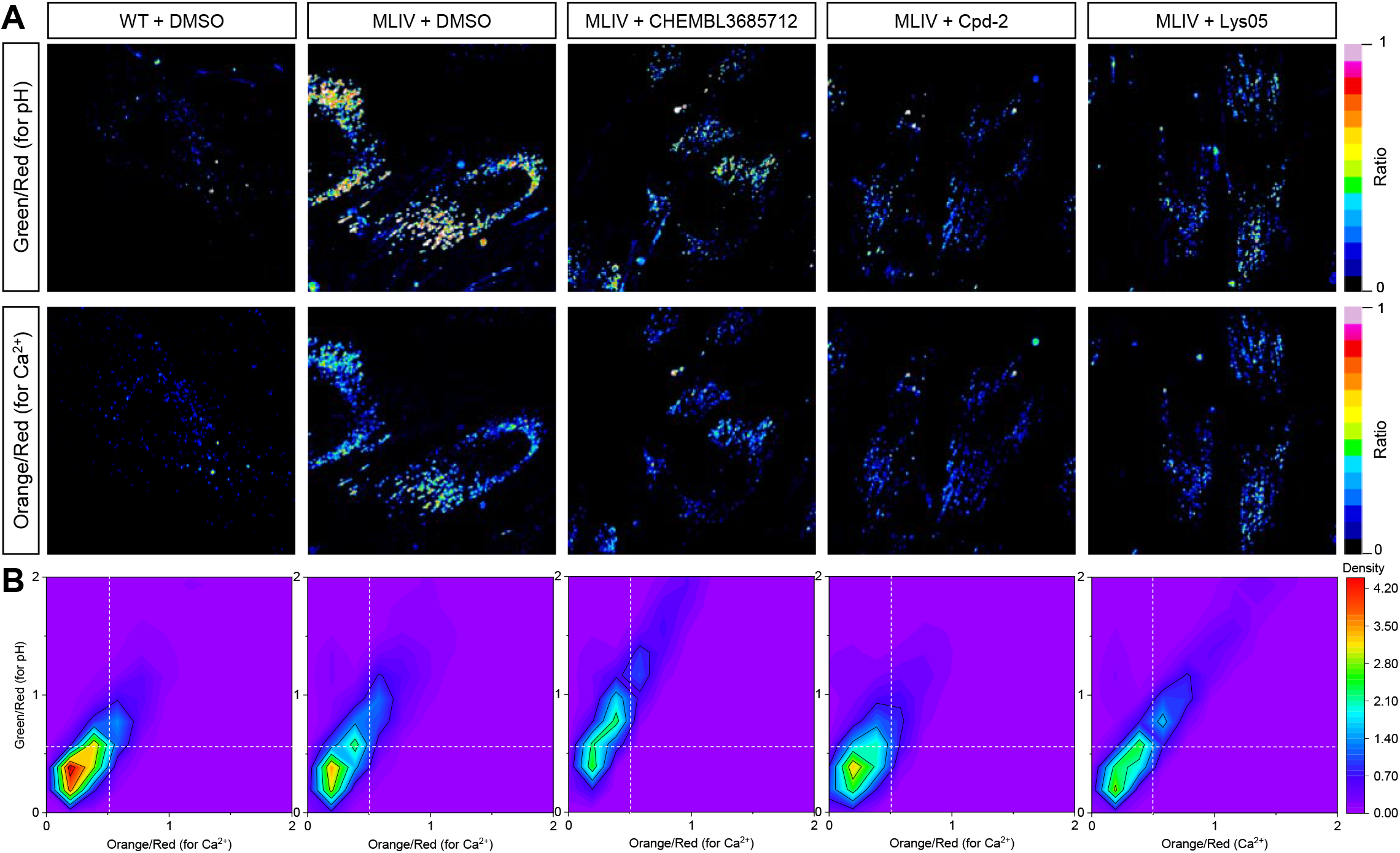
A) Representative images of CalpHlour_*Ly*_ in MLIV and wild-type fibroblast cells at *t* = 9 hours post-internalization in the presence of the indicated pharmacological inhibitors (CHEMBL3685712, Cpd-2, and Lys05) or DMSO-treated control. Representative images of acidification (Green/Red channels) in concert with [Ca^+2^] (Orange/Red channels). B) Pseudocolor image plots based on quantification of n > 1000 lysosomes in images of fibroblast cells (wild-type or MLIV) treated with DMSO or pharmacological inhibitors (CHEMBL3685712, Cpd-2, and Lys05). The y-axis represents ratio of Green/Red channels as a quantification of pH. On the x-axis is the ratio of Orange/Red channels correlating to Ca^+2^ concentration.

## Discussion

We embarked on our multi-species phenotypic screening campaign across model organisms and human fibroblasts, curious as to how a collection of well-defined chemical probes designed for genetic targets in humans would fare across invertebrate counterparts and a simplified patient-derived cell line. By focusing on the intersection of worm, flies, and human fibroblasts, we hoped we might better understand how to improve the therapeutic predictive value of model organisms. After screening the mechanism-of-action box (MoA Box) compound collection across three disease models, we did not observe overall trends that are simply explained by the *in silico* predicted bioaccumulation in *C. elegans* or relating to the shared sequence homology overlap of specific gene targets across the three species. Instead, the overlap of the screens captures similar biological pathway modulation to bypass malfunctioning TRPML1 instead of capturing other non-drug-target measures such as physiochemical (stability and solubility) or pharmacodynamic (bioaccumulation and xenobiotic detoxification) properties. Interestingly, there are only seven compounds that have activity in both an invertebrate model organism as well as the human fibroblast assay. In contrast, there are a substantially larger number of hits that are specific for *Drosophila* (86), *C. elegans* (52), or *H. sapiens* fibroblasts (167). Inclusion of the patient fibroblast assay was intended to ensure relevance to human biology and disease; however, the large number of model organism specific hits begs the question as to what mechanism or activity these compounds may be targeting in whole animals. Many of the hits that are specific to the invertebrate models (fly and worm only) are annotated for oxidative stress, nitric oxide synthesis, and DNA damage response. One possibility is that those pathways are non-specifically beneficial. Another possibility is that those pathways are indeed disease-modifying for MLIV but have no effect on LysoTracker staining in MLIV patient-derived fibroblasts. It is known that TRPML1 can act as an oxidative stress sensor and that chaperone-mediated autophagy is defective in MLIV (Zhang, et al., 2016; Venugopal, et al., 2009), but these pathways are likely less proximal to the final LysoTracker-based readout of the fibroblast assay. Pyruvate kinase M2 was an initial hit also only in flies and worms. In follow up experiments we were able to show that it has modest activity in LysoTracker clearance without observed p62 accumulation (**SI Figure 3.1**; shikonin or alkannin) (Li, et al., 2014). The observed effect may be a result of the additional antioxidant activity of the quinone scaffold (Assimopoulou, et al., 2004). Inhibition of another gene target hit, c-Abl (**SI Figure 3.2**; BAW209), showed modest reduction of LysoTracker without the induction of p62 accumulation with the exception at the highest dose, which could be due to toxicity (Sivasankaran & Zimmermann, 2009). c-Abl is a tyrosine kinase that has recently been shown to induce TFEB phosphorylation, and its inhibition can promote lysosomal biogenesis and autophagy, particularly promoting cholesterol clearance in Niemann-Pick type C models (Contreras, et al., 2020). Further studies looking at c-Abl’s role in other lysosomal storage diseases such as MLIV would be of great interest.

Most striking from our initial screens and targeted expansion compound set was the number of CDK inhibitors, which positively rescued the respective assays. CDK inhibitors have been recently shown to induce cyto-protective autophagy in cancer cells (Mathiassen, et al., 2017; Capparelli, et al., 2012; Jiang, et al., 2010; Liang, et al., 2007) and can act through phosphorylation and inactivation of Vps34 (Furuya, et al., 2010). Some studies have shown CDK inhibition to impede autophagy. For example, CDK4 inhibition impairs autophagic flux and lysosomal degradation thru modulation of mTORC1 activity (Martinez-Carreres, et al., 2019). In contrast, CDK5 inhibition has been shown to promote lysosomal biogenesis, without affecting mTOR activity (Ishii, et al., 2019). Furthermore, the gene target that was common across all three species, ILK, is structurally similar to CDK, and likewise ILK inhibits autophagy by promoting the phosphorylation of AKT and activating mTOR (Pang, et al., 2016; Sosa, et al., 2018). Other work has shown that the overexpression of ILK increases expression of various cyclins, CDK2, and CDK4 (Radeva, et al., 1997).

We were particularly interested in the role of CDK4/6 as a potential therapeutic modality for MLIV disease, as there are several approved CDK4/6 inhibitors (palbociclib and ribociclib) on the market for the treatment of ER-positive, HER2-negative metastatic breast cancer. The repurposing of CDK4/6 inhibitors would provide MLIV patients with a new option for treatment. Recent reports show that CDK4/6 can regulate autophagy and lysosomal biogenesis via the transcription factors TFEB and TFE3 (Yin, et al., 2020). In the work by Yin *et al*., they uncover a role of CDK4 and CDK6 in the nuclear export of TFEB and TFE3 through their phosphorylation, and in turn show that CDK4 and CDK6 inhibition by small molecules like palbociclib, or genetic depletion with siRNA or CRISPR knockout, activates both TFEB/TFE3 and induces lysosomal biogenesis and autophagy. This type of mechanism is consistent with rescue of the lysosomal defects in three distinct disease contexts of MLIV, where the cells of worms, flies, or MLIV patient fibroblasts, are deficient in proper autophagy because they lack a functioning cation channel TRPML1 (MCOLN1). Defective TRMPL1 leads to several abnormalities in MLIV disease, such as defects in endolysosomal and lipid trafficking, aberrant autophagy, altered homeostasis of heavy metals, and perturbations to the mTORC1/TFEB signaling axis (Boudewyn & Walkley, 2019). In wild-type cells, during starvation or low nutrient conditions, mTORC1 is inactive, and TRPML1 functions to pump calcium as well as other ions out of the lysosome. Local lysosomal calcium efflux is important for activation of calcineurin, a phosphatase that can dephosphorylate TFEB/TFE3 and allow their translocation into the nucleus to turn on lysosomal/autophagic gene expression (Boudewyn & Walkley, 2019). Without properly functioning TRPML1, even under starvation or stress conditions, TFEB/TFE3 remain phosphorylated and are unable to turn on autophagic gene expression. Our results show that in MLIV fibroblasts, we can modulate autophagy through a bypass suppressor pathway by inhibition of CDK4 and CDK6. We tested several CDK inhibitors including ribociclib, palbociclib, and non-lysosomotropic analogs of ribociclib. Through CDK inhibition, we were able to see reduction in the accumulation of highly acidic vesicles (LysoTracker signal) in MLIV cells, as well as increased levels of LC3-II levels under serum starvation conditions, indicating an increase in autophagosome/lysosomal biogenesis.

Since TRPML1 is a cation channel, with calcium being the major transported ion, we sought to look at lysosomal calcium levels using a pH-correctable Ca^2+^ reporter, CalipHlour_*Ly*_, that is directed to the lysosome (Narayanaswamy, et al., 2019). We were able to treat MLIV cells with CDK inhibitors and showed that the non-lysosomotropic analog of ribociclib, Cpd-2, was able to restore calcium and pH levels. Images of cells treated with ribociclib and BIM-1, which had very strong effects in inducing LC3-II processing, were not able to be analyzed due to mislocalization of the probe, most likely due to degradation or displacement from the lysosome.

In summary, through multi-species phenotypic screening in model organisms and patient-derived fibroblasts, we identified unpredicted drug target overlaps across *Drosophila*, *C. elegans*, and human fibroblast screens. From the trio of screens emerged a strongly suggestive role for CDK4/6 inhibition in enhancing lysosomal biogenesis in mucolipidosis Type IV models. We hope that future work will expand upon our MLIV invertebrate model organism study and evaluate FDA-approved CDK4/6 inhibitors in a mouse model of MLIV. In sum, our work has provided new insights into MLIV disease pathobiology and opens new avenues for research and potential repurposing of approved therapeutics for the MLIV patient population.

## Materials and Methods

### Compounds and strains

MK6-83 (#SML1509, Sigma-Aldrich), bafilomycin A1 (#1334, Torcis), and all MoA Box compounds were dissolved in DMSO and stored at −80°C in Labcyte-compatible LDV plates until use. *C. elegans cup-5* strains, hypomorphic strain *ar465[Pmyo-3::ssGFP]* along with wild-type *arIs37[Pmyo-3::ssGFP],* were obtained from the Fares Lab at University of Arizona, Tucson (Fares & Greenwald, 2001). *D. melanogaster* strains used for screening were generated by a CRISPR/Cas9 2 kb deletion into the trpml gene (GenetiVision). The resulting t*rpml*^*−/−*^ homozygote animals are null and have a pupal lethality/failure to eclose phenotype.

### Cells and cell culture

Mucolipidosis IV (MLIV) mutant (GM02527) and healthy wild type (GM09503) patient derived fibroblasts were purchased from the Coriell Cell Repository (Camden, NJ). Cells were cultured in EM10 media (Minimal Essential Media with GlutaMAX supplement, Gibco, # 41090-101, supplemented with 10% Fetal Bovine Serum, 1% penicillin/streptomycin, and 1% Non-Essential Amino Acids (Gibco, #11140076)) at 37°C with 5% CO_2_.

### *D. melanogaster* screen

With an Echo550 liquid handler (Labcyte Inc.), MoA box compounds (20μM) were acoustically dispensed and standard fly food media (molasses, agar, yeast, propionic acid) lacking cornmeal, but carrying 0.025% Bromophenol Blue, was dispensed using a Multi-Flo (BioTek Instruments) into each well of a 96-well plate for drug screens. Using the BioSorter (Union Biometrica) 5 1^st^ instar larvae were deposited per well and grown at 21°C. At days 13 and 14 post incubation, the plates were images using a custom built Fly Imager. The Fly Imager uses a Sony a7r ii camera, controlled over USB by the gphoto software to generate full plate images that are then run through a movement detection algorithm. At day 15 post incubation, the plates were manually analyzed to quantify the number of partially and fully hatched adult flies and inspected for cytotoxic effects. The statistical program R (https://www.r-project.org/) was used for Z-score calculations. Z-scores for test wells were calculated by normalizing by mean area s.d. of negative control wells. For each replicate, a test well that had a Z-score ≥1.5 was counted as a hit.

### *C. elegans* screen

Using the Echo550 liquid handler (Labcyte Inc.), MoA box compounds (25μM) were acoustically dispensed into the destination plates the day before the worm larvae sort. In total, 5 μl of HB101 bacteria were dispensed into 384-well plates containing S-medium (prepared in-house) in each well. Using the BioSorter (Union Biometrica), 15 L1 *cup-5* mutant larvae were sorted into each test well, and plates were incubated for 5 days at 20°C while shaking. Wild-type L1 larvae were dispensed as positive controls while untreated *cup-5* mutant L1 larvae were dispensed as negative controls into 32 wells per plate. After 5 days of benchtop incubation, 15 μl of 8 mM sodium azide was added to each well to immobilize worms prior to imaging in a custom worm imager with a 4X objective. Plates were imaged under transmitted light as well as GFP LED light source. Finally, automated image processing was run on each plate. The average area occupied by worms in positive and negative control wells as well standard deviation (s.d.) of control wells were calculated. Outlier elimination was performed by identifying those wells that were greater than 1.5 times the interquartile range (1.5× IQR). After determining the area occupied by worms, the GFP signal/ unit worm area ratio was calculated and used as the final readout. The statistical program R (https://www.r-project.org/) was used for Z-score calculations. Z-scores for test wells were calculated by normalizing by mean area s.d. of negative control wells. For each replicate, a test well that had a Z-score ≥1.5 was counted as a hit. Because image artifacts and cytotoxicity can often confound area measurements, manually verification of each well through visual analysis was done after the unbiased quantitative analysis to make sure no false positives were counted as true hits.

### MLIV patient fibroblasts cellular screen

Cells were plated at 400 cells/well in 5 microliters of media in 1536-well Aurora Ultra Low Base imaging plates (Aurora # EBC2-41001A) and cultured for 16 h at 37°C. Positive control, bafilomycin A1 (12nM), and negative control, DMSO, were included on every plate. MoA Box compounds in 8-point dose response were dispensed (Echo 555 Acoustic Dispenser) at 20 nL per well and incubated for 48 h. After incubation for 48 h, 50 nM Lysotracker Red DND-99 (LTR, Invitrogen #L7528) and 13 μM of Hoechst 33342 (Invitrogen #H3570) nuclear stain were added, incubated for 13 minutes at 37°C, and immediately imaged live on an InCell 6000 imager at 20X objective/0.75 aperture in the Red (577/591) and Blue (350/461) channels, with one field per well, in biological duplicate. For p62 immunostaining, plates were imaged live for LTR and then subsequently fixed with 4% paraformaldehyde for 20 minutes at room temperature (RT). Plates were then washed 5 times with phosphate buffered saline (PBS), and blocked for 1 h at RT using Blocking buffer (Odyssey Blocking Buffer in PBS, Licor #927-4000; 0.1% TritonX-100; 1% Goat Serum) and p62 antibody (mouse anti-p62 Ick ligand, BD Bio #610833) at 1:2000 final overnight at 4°C. Wells were washed 5 times with PBS and incubated with secondary antibody (Alexa fluor 488 goat anti-mouse IgG (H+L), Invitrogen #A11017) at 1:500 and imaged on InCell 6000 imager at 20X objective/0.75 aperture lens in the Green (488/511) and Blue (350/461) channels.

Images were then analyzed through custom pipelines generated for Cell Profiler (Stoter, et al., 2019) to measure the intensity and number of accumulated lysosomal puncta. Data was submitted to MultiParametric Data Analysis software and parameters with best statistical score for RZ’ were selected. Hit and curve calling was done by filtering to samples that had > 2 standard deviation differences in activity from the neutral control and filtered for compounds with less than 10 μM AC_50_ values. Data was also filtered for toxicity based on cell count and manual inspection of images. Immunofluorescent staining p62 acted as a filter/indication of autophagy induction/potential toxicity of compound.

Images were then analyzed through custom pipelines generated for Cell Profiler to measure the intensity and number of accumulated lysosomal puncta. Data was submitted to MultiParametric Data Analysis software and parameters with best statistical score for RZ’ were selected. Hit and curve calling was done by filtering to samples that had > 2 standard deviation differences in activity from the neutral control and filtered for compounds with less than 10 μM AC_50_ values. Data was filtered for toxicity based on cell count and manual inspection of images. Immunofluorescent staining p62 acted as a filter/indication of autophagy induction/potential toxicity of compound. **Serum Starvation for LC3B induction**

MLIV patient fibroblasts were grown for 2-3 days until 80-90% confluent and then plated in 6-well dishes in full serum media (media formulation as described above in 10% FBS) at 100,000 cells per well for 48 h starvation and 300,000 cells per well for 24 h starvation. Cells were incubated overnight before media was changed to serum starvation media (media formulation as above but with 0.1% FBS), and treated with 20 μM compounds (cells treated with Lys05 at 1 μM) for 24 or 48 h before harvesting lysates for LC3B western blots as described below.

### siRNA knockdown in patient fibroblasts

MLIV patient fibroblasts were grown for 2-3 days until 80-90% confluent at 37°C and prepared for electroporation with approximately 50nM of ON-TARGETplus siRNA SMARTpool reagents from Horizon Cell Solutions (Dharmacon) according to manufacturer instructions using the Amaxa P2 Primary Cell 4D-Nucleofector X Kit (Lonza, Cat. No. V4XP-2032) with nucleocuvette strips in a Lonza 4D-Nucleofector X Unit. In brief, cells were washed with 30 mL PBS, and lifted for 3 min at 37°C using 5 mL of TrypLE Express (Invitrogen, Gibco, Cat no. #1260501). The appropriate number of cells were then spun down at 90 x g for 10 min at room temperature, for final 100,000 cells per siRNA condition. Cells were resuspended in 20 μl of prepared nucleofector solution with added supplement, and 2 μl of siRNA solution (final 50 nM of siRNA only, or siRNA and GFP plasmid control (0.4 μg)) was added to each tube containing cells. Cell/siRNA solution was then transferred to nucleocuvette strips, and gently tapped to remove any bubbles. Nucleocuvette strip was then placed in the Lonza 4D-Nucleofector X Unit (Lonza, Cat No. AAF-1002X) for electroporation using program DT-130 for NHDF P2 Primary cells. After nucleofection, samples were incubated at room temperature for 10 min before being added to pre-warmed media in plates for imaging or western blots. Cells were then grown for 72 hrs for imaging assays or 5 days for western blot protein analysis. For imaging analysis, cells were treated following protocol above for Lysotracker live staining and p62/TFE3 fixed staining.

siRNAs used included ON-TARGETplus SMARTpool siRNA Human Reagents from Horizon Cell Solutions (Dharmacon) for ILK (Entrezgene 3611; Cat. No. # L-004499-00-0005), CDK2 (Entrezgene 1017; Cat. No. #L-003236-00-0005), CDK4 (Entrezgene 1019; Cat. No. #L-003238-00-0005), CDK5 (Entrezgene 1020; Cat. No. #L-003239-00-0005), CDK6 (Entrezgene 1021; Cat. No. #L-003240-00-0005), CDK9 (Entrezgene 1025; Cat. No. #L-003243-00-0005), GBA (Entrezgene 2629; Cat. No. #L-006366-00-0020), and Non-targeting (NT) Control siRNA #1 (Cat. No. #D-001810-01-20).

### Western Blot Protein Analysis

Cells were harvested and lysates prepared for western blot by addition of 100 μl RIPA buffer containing EDTA, EGTA (Boston Bioproducts, Cat. No. #BP-115DG) and Protease/Phosphatase Inhibitors (Halt, Thermo Cat. No. #78442). Lysates were kept on ice and vortexed 4 times over 20 minutes, and cleared by centrifugation at 14000 rpm for 10 min at 4°C. Lysate protein concentration was determined using the Direct Detect Spectrometer (Millipore Sigma, Cat. No. #DDHW00010-WW) and normalized to 5 μg. Samples were prepared with 1X LDS Sample loading buffer, and boiled for 10 min at 70°C. Proteins were then separated using gel electrophoresis using 4-12% Bis-Tris Novex Midi 20 well gels (Invitrogen, Cat. No. #WG1402BOX) with 1X MES Buffer (Invitrogen, Cat. No. #NP0002) and run at 130V for 1.5 h. Proteins were then transferred using Bio-Rad Transblot system for Semi-dry transfer (BioRad Midi, Cat. No. #1704159) for 7 min at 2.5A/25V. Membranes were then blocked by incubation in 3% milk (Cat no.) in Tris-Buffered Saline with 0.1% Tween-20 (TBS-T) for 1 h at RT, before addition of primary antibody for incubation at 4°C overnight. Antibodies used included LC3B (Rabbit, Cell Signal, 2775S) at 1:1000 in 5% Bovine Serum Albumin (BSA) in TBS-T, p62 (Mouse, BD Biosciences, 610833) at 1:3000 in 5% BSA in TBS-T, and Actin (Mouse, Sigma, A5441) at 1:100,000 in 3% milk in TBS-T. After overnight incubation, membranes were washed 5 times with TBS-T and incubated with secondary antibodies in 3% milk in TBS-T (Anti-Rabbit IgG HRP linked, Cell Signal, #7074S; Anti-Mouse IgG HRP linked, Cell Signal, #7076) for 1 h at RT. Membranes were then washed 5 times with TBS-T before incubation with ECL substrate for 60 seconds (SuperSignal West Dura Extended Duration Chemiluminescent ECL substrate, 1:1, Thermo Scientific, Cat. No. 34075 or SuperSignal West Femto Maximum Sensitivity ECL Substrate, 1:1, Thermo Scientific, Cat. No. #34095). Blots were exposed and developed using film (GE Amersham Hyperfilm ECL, Cat. No. 28-9068-35) for intervals of 10 sec to 5 min. For quantification of Western blots, bands were quantified using Image J and normalized to protein loading controls (Schneider, et al., 2012). Statistical analysis was performed using a Student’s unpaired two-tailed t-test.

### Calcium Imaging using CalipHlour_Ly_

MLIV and wild type fibroblasts were plated at 400 cells/well in 5 microliters of media in 1536-well Aurora Ultra Low Base imaging plates (Aurora # EBC2-41001A) and cultured for 16 h at 37°C. Compounds were added at 20 μM final concentration using Echo 555 Acoustic Dispenser and incubated 48 h. Aqueous CalipHlour_*Ly*_ probe (Esya Labs, Inc.; esyalabs.com) was dispensed using the Echo Dispenser at 500 nM final concentration, and incubated for 9 h. Plates were then washed two times with Hanks Balanced Salt Solution (HBSS) and 5 μM final Hoechst 33342 nuclear stain was added. For image/microscope calibration, CalipHlour_*Ly*_ coated beads (Esya Labs, Inc; esyalabs.com) were washed with binding buffer (sodium acetate, pH 4.6), and then incubated in Low Calcium or High Calcium Buffers. Beads were then diluted 1:100 and 5 microliters were placed in empty wells of 1536-well plate containing cells. Cells were then imaged live at 40X using Yokogawa CV8000 microscope. Images were collected in four channels – Blue (350/461), Green (488/511), Orange (571/591) and Far Red (647/667) and image analysis for quantification of pH and calcium was conducted as described in (Narayanaswamy, et al., 2019).

## Acknowledgements

We thank Johnny Fares (U of Arizona, Tucson) for the *cup-5 C. elegans* strains *ar465[Pmyo-3::ssGFP]* and *arIs37[Pmyo-3::ssGFP]*. We acknowledge the team from the Astex-Novartis collaboration for their work on the development of ribociclib and Cmpd-2. We gratefully thank Walter Michaels and Christopher Brain (Novartis, Cambridge, MA USA) for a sample of Cmpd-2. We thank Dhivya Venkat (Esya Labs) for help in acquisition of the CalipHlour_Ly_ probe. We appreciate the help of Souvik Modi (Esya Labs) and Yamuna Krishnan (University of Chicago & Esya Labs) for help in experimental design and data analysis and interpretation with CalipHlour_*Ly*_ probe. We would like to acknowledge the help of Avi Spier and Risa Shapiro (Novartis, Cambridge, MA USA) for their help in this collaborative research project.

## Author contributions

A.H., S.I., J.D.M., N.D. designed and led the study under the supervision of E.O.P., N.T.R., R.K.J., J.A.T., and S.M.C. A.H., S.I., J.D.M., N.D., J.K., T.P.R., F.S.S., H.T., M.P., J.B., K.M., T.A.H. developed the methodology and carried out the experimental investigations detailed in this study. A.H., S.I., J.D.M, N.D., H.T., K.M., J.K., F.S., S.M., Y.K., C.A., B.N., E.O.P, and S.M.C. analyzed results and data generated from the study. J.L.J. determined the *in silico* bioaccumulation score for the Novartis MoA Box. S.M.C., E.O.P., J.A.T., R.K.J., N.T.R. conceptualized and supervised the study. A.H. and S.M.C. prepared the initial draft of the manuscript with input from all of the authors.

